# Targets and cross-reactivity of human T cell recognition of Common Cold Coronaviruses

**DOI:** 10.1101/2023.01.04.522794

**Authors:** Alison Tarke, Yun Zhang, Nils Methot, Tara M. Narowski, Elizabeth Phillips, Simon Mallal, April Frazier, Gilberto Filaci, Daniela Weiskopf, Jennifer M. Dan, Lakshmanane Premkumar, Richard H. Scheuermann, Alessandro Sette, Alba Grifoni

## Abstract

The Coronavirus (CoV) family includes a variety of viruses able to infect humans. Endemic CoVs that can cause common cold belong to the alphaCoV and betaCoV genera, with the betaCoV genus also containing subgenera with zoonotic and pandemic concern, including sarbecoCoV (SARS-CoV and SARS-CoV-2) and merbecoCoV (MERS-CoV). It is therefore warranted to explore pan-CoV vaccine concepts, to provide adaptive immune protection against new potential CoV outbreaks, particularly in the context of betaCoV sub lineages. To explore the feasibility of eliciting CD4^+^ T cell responses widely cross-recognizing different CoVs, we utilized samples collected pre-pandemic to systematically analyze T cell reactivity against representative alpha (NL63) and beta (OC43) common cold CoVs (CCC). Similar to previous findings on SARS-CoV-2, the S, N, M, and nsp3 antigens were immunodominant for both viruses while nsp2 and nsp12 were immunodominant for NL63 and OC43, respectively. We next performed a comprehensive T cell epitope screen, identifying 78 OC43 and 87 NL63-specific epitopes. For a selected subset of 18 epitopes, we experimentally assessed the T cell capability to cross-recognize sequences from representative viruses belonging to alphaCoV, sarbecoCoV, and beta-non-sarbecoCoV groups. We found general conservation within the alpha and beta groups, with cross-reactivity experimentally detected in 89% of the instances associated with sequence conservation of >67%. However, despite sequence conservation, limited cross-reactivity was observed in the case of sarbecoCoV (50% of instances), indicating that previous CoV exposure to viruses phylogenetically closer to this subgenera is a contributing factor in determining cross-reactivity. Overall, these results provided critical insights in the development of future pan-CoV vaccines.

## INTRODUCTION

Coronaviruses (CoV) remain a general concern because of their pandemic potential, as illustrated not only by the SARS-CoV-2 pandemic but also by previous CoV outbreaks, including SARS-CoV and the Middle East respiratory syndrome coronavirus (MERS-CoV). In this context, the development of a pan-CoV vaccine to preemptively provide adaptive immunity against the threat of a new CoV outbreak resulting from zoonotic spillover from an animal reservoir species into humans is of interest^1^. Indeed, all CoVs associated with the recent outbreaks had zoonotic origins, infecting bats, pigs, pangolins and rodents before being transferred to humans. Zoonotic and human coronaviruses, are broadly classified in two main genera: alpha and beta. The beta coronaviruses are subdivided into additional subgenera, with the beta sarbecoCoV group being of the most pandemic concern^2^. Compared to alphaCoV, the betaCoV genus has been evolutionary more prolific with multiple subgenera infecting humans with various degrees of phylogenetic relation, including merbecoCoV (MERS-CoV) and sarbecoCoV (SARS and SARS-CoV-2)^2–4^. Within alphaCoV, two CoVs infect humans seasonally (HCoV-229E and HCoV-NL63), and within the betaCoV, HCoV-HKU1 and HCoV-OC43 cause common cold upon infections, with cyclical and alternating patterns of prevalence in different populations and geographical locations^5^. These four common cold coronaviruses (CCC) are prevalent worldwide and usually cause mild illness primarily affecting the upper respiratory tract^5^.

Despite the seasonality and prevalence of common cold viruses, scarce data were available regarding general immunity, with a prevalent focus on humoral immunity and no information on cellular immunity before the pandemic ^6^. In contrast, both components of adaptive immunity have been more investigated in the context of the sarbecoCoV, with particular emphasis on SARS-CoV-2^1,7,8^.

In the context of SARS-CoV-2 and with particular emphasis on cellular immunity, several studies have shown that early broad and polyantigenic T cell responses play a potential part in the resolution of SARS-CoV-2 infection and COVID-19 ^7,9–11^. In non-human primates, T cells can also contribute to the reduction of SARS-CoV-2 viral loads ^12^. When individuals with agammaglobulinemia and B cell-depletion are infected with SARS-CoV-2, there is only a small increase in the risk of hospitalization ^13^ indicating the T cells could be providing protection against more severe disease. Indeed, individuals with multiple sclerosis (MS) who are without antibody responses while treated with ocrelizumab, exhibit mild COVID-19 upon infection, suggesting that COVID-19 can be modulated without antibody responses ^14,15^. Furthermore, the preservation of T cell reactivity against variants in which binding of neutralizing antibodies is impaired correlates with the preservation of protection against severe disease, but decreased protection from infection. Specifically, several studies showed that T cell responses were largely preserved at the population level to SARS-CoV-2 variants, including Omicron ^16–22^.

The impact of cross-reactive T cells recognizing viral variants has also been reported in the influenza system. It has been shown that pre-existing T cell immunity to influenza correlates with disease protection ^23,24^ including protection from symptomatic infection ^25^. Additionally, influenza-specific T cell epitopes have been identified and shown to be recognized by multiple donors and conserved in multiple influenza strains ^26^. In the absence of cross-reactive neutralizing antibodies, conserved virus-specific T cell responses correlated with cross-protection from symptomatic influenza ^27^. This evidence points to the potential value of crossreactive T cells in the context of influenza viral infection and that could be applicable to other viral infections including coronaviruses.

In this context, several studies have reported pre-existing cross-reactive memory T cells associated with CCC and pre-existing T cells were shown to associate with milder disease and better vaccination responses ^7,28–32^. Specifically, T cell responses to SARS-CoV-2 have been identified in unexposed subjects ^7,28,33–35^, which in some instances, have been shown to map to cross-reactive recognition of the SARS-CoV-2 sequences by T cells induced by endemic CCC ^36–39^. SARS-CoV-2-specific T cells were also able to cross-recognize other human CoVs, including SARS-CoV-1 and MERS-CoV ^40–44^ and other viral species ^29,45,46^. Based on these data it has been proposed that immunodominant T cell regions conserved across CoVs may be of interest for inducing a panCoV T cell response ^1^, to be considered not as an alternative but in conjunction with the induction of broadly reactive antibody responses ^47,48^.

However, while over 100 different studies have investigated the T cell epitope repertoire induced after infection with SARS-CoV-2, as reviewed by Grifoni et al. in 2021 ^49^, sparse data are available for other human circulating CoVs. We have recently shown that CCC T cell immunity is readily detectable in the general population with unknown CCC exposure, and showed that it is sustained over time ^50^. This suggested the feasibility of the study of the pattern of protein immunodominance and T cell epitope repertoire recognized by the general population after CCC exposure. Accordingly, in this study, we define CCC CD4^+^ T cell targets using NL63 and OC43 as prototypes for alpha and beta CCC viruses, to study which antigens and epitopes are recognized and to what extent broad T cell responses can be identified and predicted on the basis of sequence conservation.

## RESULTS

### Characteristics of a cohort of healthy blood donors who donated pre-SARS-CoV-2 pandemic samples

We investigated the pattern of immunodominance in T cell responses using PBMC isolated from 88 healthy adult participants (**Table 1;**indicated hereafter as “First Cohort”), spanning a wide age range (ages from 19 to 84 years; median 46), with a balanced sex ratio (48:52, Male:Female). The ethnic breakdown was reflective of the demographics of the local enrolled population with a prevalence of white-not Hispanic or Latino (60%), and a 22% representation of other ethnicities, 18% of the cohort has not reported information regarding ethnicity. HLA typing of these donors is presented in **Table S1**. Blood samples were collected from March 2020 to February 2021. Accordingly, and based on epidemiological data on CCC seasonality for the 2019-2021 years ^5^, we selected NL63 and OC43 as representative prototypes for recent exposure to alphaCoV and betaCoV CCC, respectively.

**Table 1.** Summary of the cohort analyzed in this study.

|  | First Cohort<br>(n = 88) | Validation Cohort<br>(n = 7) |
| --- | --- | --- |
| **Age (years)** | 19-84 [Median = 45, IQR = 30] | 23-74 [Median = 59, IQR = 46] |
| **Sex** |  |  |
| Male (%) | 48% (43/88) | 86% (6/7) |
| Female (%) | 52% (46/88) | 14% (1/7) |
| **Sample Collection Date** | March 2020 – February 2021 | October 2018 – August 2019 |
| **Race-Ethnicity** |  |  |
| White- not Hispanic or Latino | 60% (53/88) | 44% (3/7) |
| Hispanic or Latino | 15% (13/88) | 14% (1/7) |
| Asian | 6% (5/88) | 14% (1/7) |
| Black or African American | 1% (1/88) | 14% (1/7) |
| Not reported | 18% (16/88) | 14% (1/7) |

All donors in this cohort were seronegative for SARS-CoV-2 Spike RBD antibodies at the time of sample collection, ensuring that any responses detected would not be related to SARS-CoV-2 exposure or vaccination. Likewise, as expected based on expected previous CCC exposure, these donors were seropositive for antibodies to the RBD domains of the NL63 and OC43 viruses, as measured by IgG ELISA (**Fig. S1**).

### Immunodominant antigens recognized by CCC-specific T cell responses

In contrast with the wealth of information available on antigen immunodominance resulting from SARS-CoV-2 infection ^49^, a systematic analysis of which antigens are dominantly recognized in T cell responses elicited by CCC infection is currently lacking. To define the specific antigens recognized by CD4^+^ T cells from donors previously exposed to human alpha and beta coronaviruses (as determined by NL63 and OC43 seropositivity), we tested PBMCs from the First Cohort donors described in **Table 1**, with sets of overlapping peptides spanning proteins from the entire viral proteome (**Fig. 1**). The same approach was previously used to define the SARS-CoV-2 antigens recognized by T cells^51,52^ in SARS-CoV-2 convalescent donors. CD4^+^ T cell responses were measured by the Activation Induced Marker (AIM) assay utilizing the OX40 and CD137 markers. The gating strategy of the flow cytometry-based AIM assay is detailed in **Fig. S2A**. For each antigen/donor combination, the total magnitude of response is shown as a heatmap to illustrate appreciate the donor-to-donor variability (**Fig. 1A-B**). Additionally, for each protein, both the total magnitude, calculated by summing all responses observed for a given antigen in the study cohort, and frequency of responses were derived (**Fig. 1C-D**).

**Figure 1.**
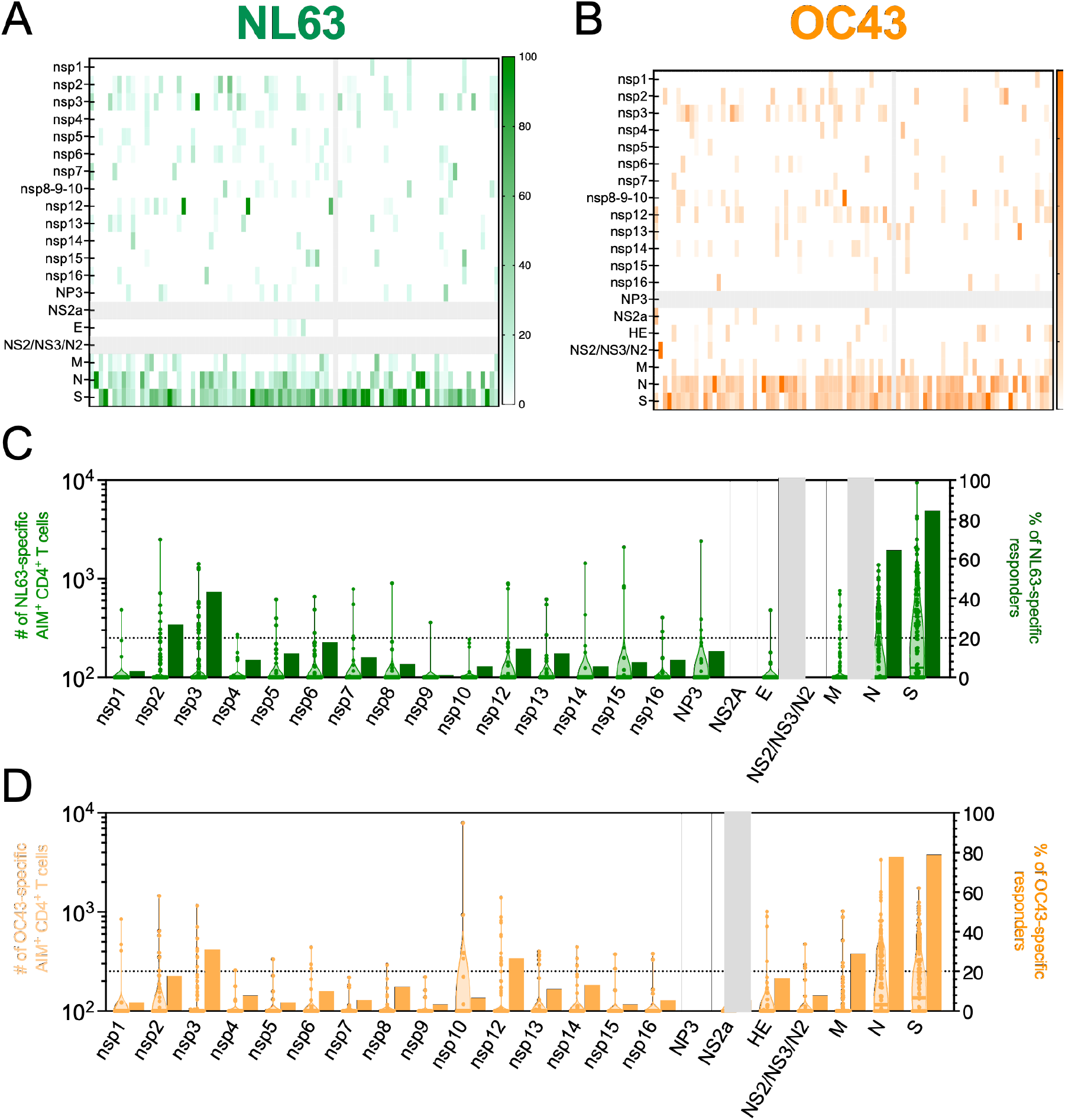
CCC-specific CD4^+^ T cell reactivity per protein. PBMCs from healthy donors (n=88) were analyzed for reactivity against NL63 (green; **A** and **C**) and OC43 (orange; **B** and **D**). **A-B**) T cell reactivity across the CCC proteome is shown as heatmaps as a function of the donor tested. The x-axis shows individual donors’ responses for each protein (y-axis). **C-D**) Immunodominance at the antigen level and for the frequency of T cell responders. Magnitude data per each single donor/protein combination are shown as truncated violin plots on the left y-axis. The frequency of donors responding to the specific protein are shown as bar plots on the right y-axis.

The overall pattern of recognition of NL63 and OC43 was similar, specifically, 20% or more of responses were ascribed to the S, N, M, or nsp3 proteins for both viruses, with S and N proteins most dominantly recognized, followed by M and other non-structural proteins (**Fig. 1C-D**). Additionally, nsp2 and nsp12 responses were found to be more frequent in NL63 (**Fig. 1C**) or OC43 (**Fig. 1D**), respectively. These six proteins account for 85% and 81%of the overall responses for NL63 and OC43, respectively (data not shown).

The protein antigens found here to be dominant in CCC responses, were similar to those previously shown to be dominantly recognized in the context of SARS-CoV-2 responses ^52^. In the case of CCC responses, the top 6 antigens accounted for 80% or more of the Alpha and Beta non-SARS-CoV-2 responses, as compared to the 8-9 protein antigens required to cover 80% of the SARS-CoV-2-specific response ^52^.

To assess relative dominance hierarchies, we next plotted the total T cell reactivity detected for each OC43 and NL63 antigen in the current study, and compared this to the total reactivity to the various SARS-CoV-2 antigens, previously measured in a cohort of mostly mild COVID-19 convalescent donors using the same methodology ^52^. The majority of the antigens were similarly recognized in the different viruses, with Spearman correlation analysis (**Fig. S2B**) showing significant positive correlations, especially for OC43/SARS-CoV-2 (R=0.5649 and p=0.0095) and NL63/SARS-CoV-2 (R=0.6140 and p=0.0067). This is consistent with those two viruses belonging to the betaCoV genus, and therefore being phylogenetically more similar to each other than with the NL63, which belongs to the alphaCoV genus ^53^.

The total SARS-CoV-2 response previously observed in SARS-CoV-2 convalescents^52^ was significantly higher than what was observed with the other two CCC (Kruskal Wallis; P<0.0001), consistent with the more recent SARS-CoV-2 exposure in the COVID-19 convalescent cohort, which was analyzed 1 month after infection, as compared to the unknown timing of exposure to the other CCC in the donors analyzed in this study (**Fig. S2C**). Overall, these results provide the first unbiased genome wide analysis of CD4^+^ T cell reactivity to two ubiquitous CCC.

### Identification of CCC-derived CD4^+^ T cell epitopes

The analysis of the NL63 and OC43 proteome-wide immunogenicity pinpointed specific donor-antigen combinations associated with good reactivity for CD4^+^ T cells, that were deconvoluted based on the availability of donor cells. To identify specific CD4^+^ T cell epitopes, we deconvoluted peptide pools corresponding to the six immunodominant antigens (S, M, N, nsp2, nsp3 and nsp12) identified above as accounting for 80% or more of the NL63 and OC43 CD4^+^ T cell activity (**Fig. 1**). Epitope deconvolution was performed in at least 8 independent donors per antigen. CD4^+^ T cell epitopes were defined using an HLA-unbiased approach. First, overlapping peptides spanning the entire sequence of the antigen in question were pooled in intermediate pools of about 10 peptides each, and tested for reactivity in the AIM assay. The intermediate pools found to be positive in the AIM assays, were then deconvoluted to identify the specific peptides associated with the positive response in a second round of experiments ^52^. The positivity threshold was defined as >100 net AIM^+^ cell counts (background subtracted by the average of triplicate negative controls) and a Stimulation Index (SI) >2, as previously described ^52,54^.

**Table S2** provides a summary of the 165 epitopes identified, fairly evenly distributed between NL63 (n=87) and OC43 (n=78). Overall, these results provide the first unbiased genome-wide CD4^+^ T cell epitope identification screen to two CCC.

### Selection of a panel of alpha, beta and sarbeco virus representatives of coronaviruses of human concern

We next addressed to what degree the CCC epitopes identified in this study were conserved within different coronavirus species, with the ultimate goal of identifying CCC-specific epitopes that would be predicted to cross-react with other alpha and beta-coronaviruses, and potentially broadly reactive with other different HCoVs, including SarbecoCoV of potentially pandemic concern, as well. Several studies have reported pre-existing memory SARS-CoV-2 CD4^+^ T cell reactivity in unexposed donors and have shown crossreactivity between SARS-CoV-2 and CCC sequences^35,36,39,51^, demonstrating the presence of T cell memory clones able to cross-recognize multiple HCoVs.

We selected a representative set of viruses according to the criteria summarized in **Fig. S3**. The selection included clustering based on genomic sequence identity, sorting clusters based on cluster size, and a phylogeny and metadata-based sampling method to select representatives (blue) in major phylogenetic clusters. A total of 33 sequences were selected including the prototype sequences used to identity NL63 and OC43 epitopes. Of those, 16 sequences were selected to represent the alphaCoV genus and 17 to represent betaCoV genus. The betaCoV were further divided into a group of 4 sequences specifically related to the subgenus sarbecoCoV and a group of 13 non-sarbecoCoV (**Table 2**).

**Table 2.**
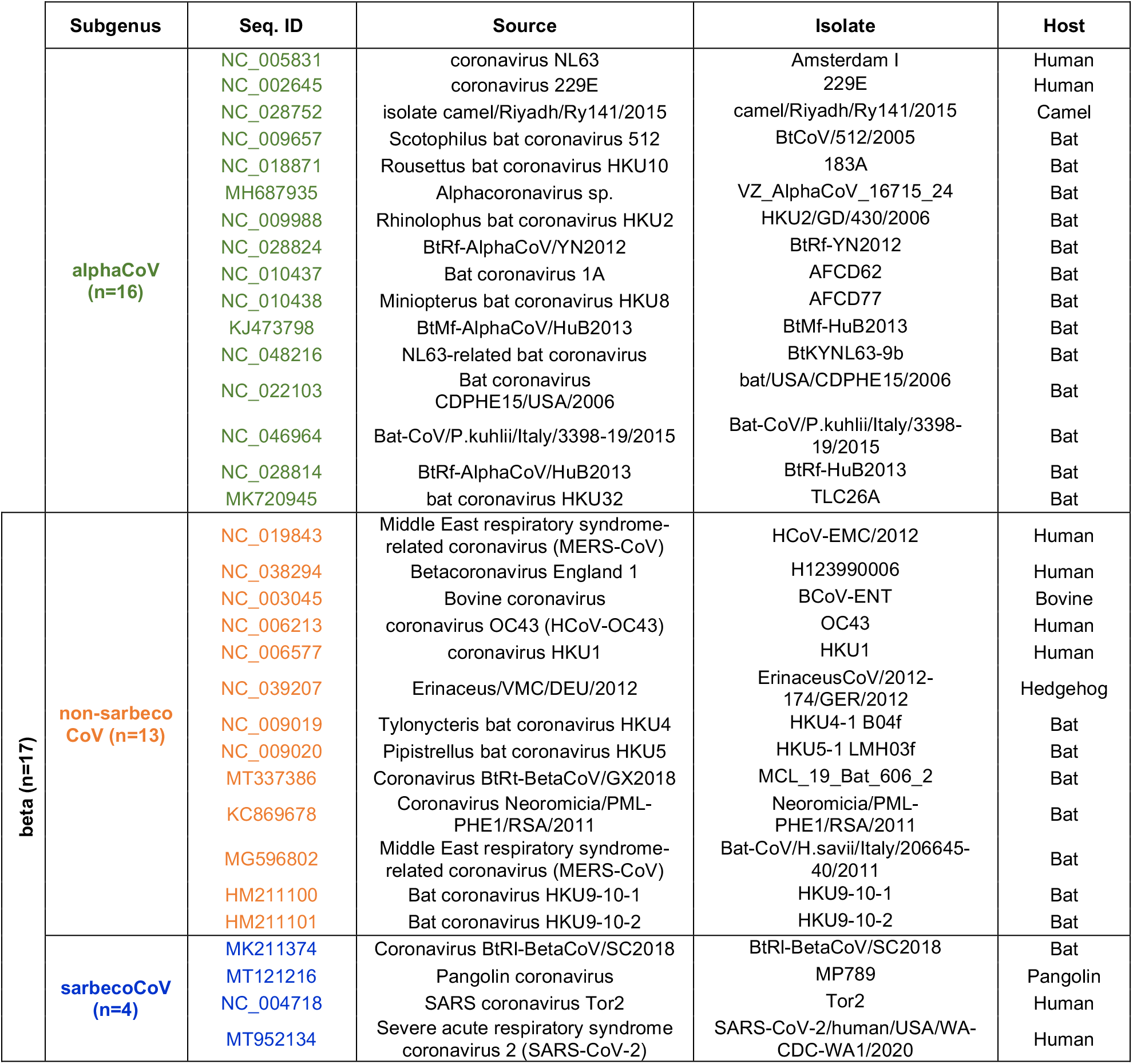
List of the representative CoVs.

### Sequence conservation of CCC CD4^+^T cell epitopes in other Coronaviruses

We first calculated the degree of conservation (% sequence conservation) of the sequences of each of the NL63 and OC43 T cell epitopes in each of the representative sequences (**Figure S4)**. We next calculated the median overall conservation for each of the epitopes, and the median conservation within alphaCoV, sarbeco, and non-sarbeco betaCoV groups (**Table S2)**. The results are also graphically summarized in **Figure 2**. Information regarding the protein location of each epitope and the total magnitude of responses associated with each epitope is shown in the vertical line graph.

**Figure 2.**
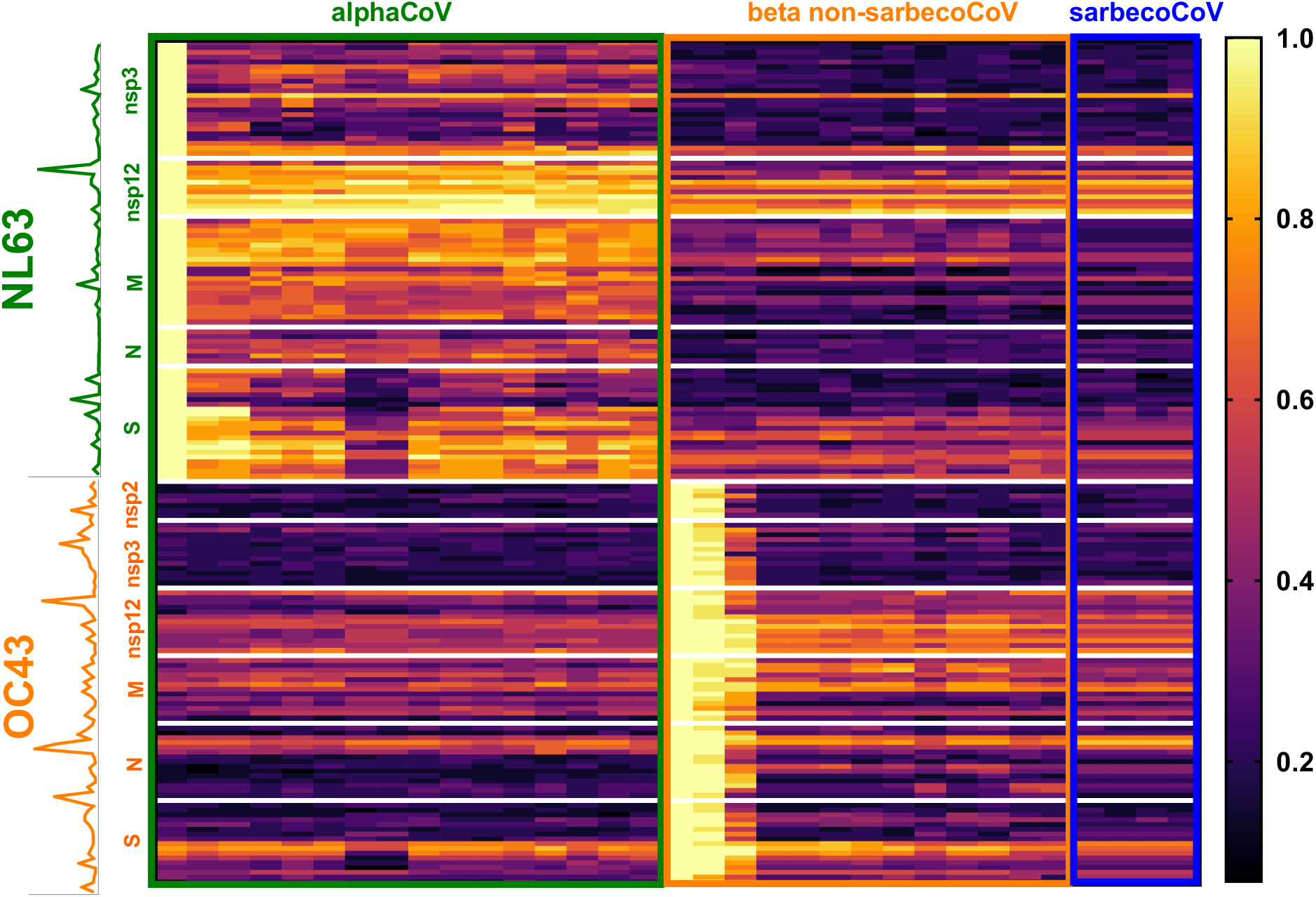
Epitope conservation across coronaviruses. Heatmap of conservation of NL63 (green text) and OC43 (orange text) T cell epitopes in each of the coronavirus representative sequences divided into alphaCoV (n=16; green outline), betaCoV non-sarbecoCoV (n=13; orange outline) and sarbecoCoV (n=4; blue outline) groups. Each row represents a different epitope, and each column a different coronavirus representative. The color intensity, following the gradient shown on the right of the heatmap, shows the degree of calculated sequence conservation for each epitope/virus combination. The y-axis shows information regarding the protein of each epitope and a line graph illustrates the total response associated with each epitope.

Overall, NL63 epitopes showed the highest degree of conservation across alphaCoV representative sequences, with some specific epitopes, especially in the nsp12 protein, associated with broad conservation across all HCoVs. OC43 epitopes show the broadest conservation particularly for nsp12, followed by specific sub-regions pertaining to M, N and S proteins. Thus, the nsp12 protein is associated with the highest combined level of conservation and immunogenicity, for both CCC viruses analyzed, making it a potential candidate to stimulate broadly cross-reactive T cell responses. In summary, the combined experiments identify a number of different epitopes and antigen regions, derived from CCC, immunogenic in humans, and with different degrees of conservation in coronaviruses of human concern.

### Selection of a panel of CCC T cell epitopes to investigate cross-recognition within other Coronaviruses

We next experimentally investigated whether T cells specific for CCC epitopes could cross-recognize peptides corresponding to the different representative coronaviruses described above. In addition to providing direct evidence for cross-reactivity within CoVs, this set of experiments was designed to determine how frequently cross-reactive recognition by human memory T cells of coronavirus sequences could be observed, and which level of homology would correlate with cross-reactivity. Previous studies had indicated a level of conservation of 67% (0.67) as being associated with cross-reactivity of SARS-CoV-2 reactive T cells with CCC sequences ^36^.

To address these points, we calculated the median conservation for each of the three groups analyzed (alphaCoVs, beta non-sarbecoCoV, and sarbecoCoV) and selected for analysis eighteen representative epitopes, defined as the sequence for which reactivity was detected in *ex vivo* experiments, associated with different degrees of conservation in the different coronavirus groups (alpha, beta non-sarbeco and sarbeco). **Table 3** details the epitopes selected, with the first column reporting the virus species from which the epitope was identified, and the specific protein and residues. The next columns describe the median conservation in the different viral groups. More specifically, 11 of the 18 representative epitopes were conserved within alphaCoV, 9 epitopes were conserved within beta non-sarbecoCoV, and 8 epitopes were conserved within sarbecoCoV with a sequence identity >0.67 (**Table 3**). Based on their pattern of conservation, as indicated by the next column in **Table 3**, 5 epitopes were classified as “common,” that is conserved in all three viral groups, 5 epitopes were conserved only in alphaCoV, 2 conserved only in betaCoV (not conserved in alpha, but conserved in all betaCoV), 2 peptides were conserved only in betaCoV-non-sarbeco, and 1 peptide was conserved only in sarbeco viruses. The last 2 peptides had sequence identities of <0.67 and were not well conserved in any of these groups.

**Table 3.** List of representative epitope candidates and experimental outcome of T cell lines.

| CCC | Epitope | % Conservation |  |  |  | TCL reactivity |  |  |  |
| --- | --- | --- | --- | --- | --- | --- | --- | --- | --- |
|  |  | alphaCoV | beta non-sarbecoCoV | sarbecoCoV | conservation category | First Cohort |  | Validation Cohort |  |
|  |  |  |  |  |  | alpha/beta reactivity | Sarbeco reactivity | alpha/beta reactivity | Sarbeco reactivity |
| NL63 | nsp12<sub>4476</sub> | 0.87 | 0.73 | 0.73 | common | TCL-only | - | n.d. | n.d. |
| NL63 | nsp12<sub>4896</sub> | 1 | 0.87 | 0.87 | common | alpha/beta | yes | n.d. | n.d. |
| OC43 | N<sub>121</sub> | 0.67 | 0.8 | 0.87 | common | alpha/beta | - | alpha/beta | yes |
| NL63 | nsp3<sub>1286</sub> | 0.87 | 0.73 | 0.8 | common | alpha/beta | - | n.d. | n.d. |
| OC43 | S<sub>911</sub> | 0.67 | 0.73 | 0.73 | common | alpha/beta | yes | alpha/beta | yes |
| NL63 | S<sub>956</sub> | 0.87 | 0.47 | 0.33 | alpha | alpha | - | n.d. | n.d. |
| NL63 | S<sub>951</sub> | 0.87 | 0.27 | 0.13 | alpha | alpha | - | alpha/beta | - |
| NL63 | nsp12<sub>4136</sub> | 0.87 | 0.33 | 0.27 | alpha | alpha | - | n.d. | n.d. |
| NL63 | N<sub>121</sub> | 0.73 | 0.53 | 0.6 | alpha | n.d. | n.d. | alpha/beta | yes |
| NL63 | M<sub>111</sub> | 0.73 | 0.47 | 0.47 | alpha | n.d. | n.d. | alpha | - |
| NL63 | nsp12<sub>4731</sub> | 0.8 | 0.33 | 0.4 | alpha | n.d. | n.d. | alpha | yes |
| OC43 | N<sub>116</sub> | 0.47 | 0.73 | 0.73 | beta | beta | yes | beta | yes |
| OC43 | nsp12<sub>4731</sub> | 0.47 | 0.73 | 0.73 | beta | beta | - | - | n.d. |
| OC43 | M<sub>36</sub> | 0.33 | 0.73 | 0.4 | non-sarbeco | alpha | - | n.d. | n.d. |
| OC43 | M<sub>46</sub> | 0.47 | 0.73 | 0.53 | non-sarbeco | n.d. | n.d. | beta | - |
| OC43 | N<sub>126</sub> | 0.53 | 0.53 | 0.67 | sarbeco | alpha | - | n.d. | n.d. |
| OC43 | S<sub>951</sub> | 0.33 | 0.4 | 0.47 | none | n.d. | n.d. | beta | yes |
| OC43 | S<sub>956</sub> | 0.33 | 0.6 | 0.6 | none | n.d. | n.d. | beta | yes |

"n.d." = not determined, indicates not tested or lack of response of the TCL; "-" = no T cell reactivity

### Assessment of cross-reactivity patterns of CCC CD4^+^T cell epitopes within other Coronaviruses

Next, we determined the pattern of cross-reactivity of T cells recognizing the various epitopes, by generating epitope-specific short-term T cell lines (TCL) from PBMCs of a subset of donors from the First Cohort. This first round of experiments investigated the reactivity of TCLs specific for 12 different epitopes. These TCL were tested with a dose range of synthetic peptides corresponding to the sequence of the homolog peptides from each of the virus isolates from **Table 2,**as previously reported ^36^. The specific TCL reactivity for each epitope and virus sequence is shown in **Figure 3A-E**. Reactivity against alphaCoV sequences is shown in green, beta non-sarbecoCoV in orange, and sarbecoCoV in blue. The reactivity of epitopes conserved in all three groups is shown in **Fig. 3A**, alpha-specific (conserved only in alpha-corona) in **Fig. 3B**, beta-specific (conserved only in beta-corona) in **Fig. 3C**, beta non-sarbeco-specific in **Fig. 3D**, and sarbeco-specific in **Fig. 3E**. The results are also summarized as heatmaps in **Figure 3F**,depicting the epitope conservation across each viral species and the reactivity of each TCL against each peptide.

**Figure 3.**
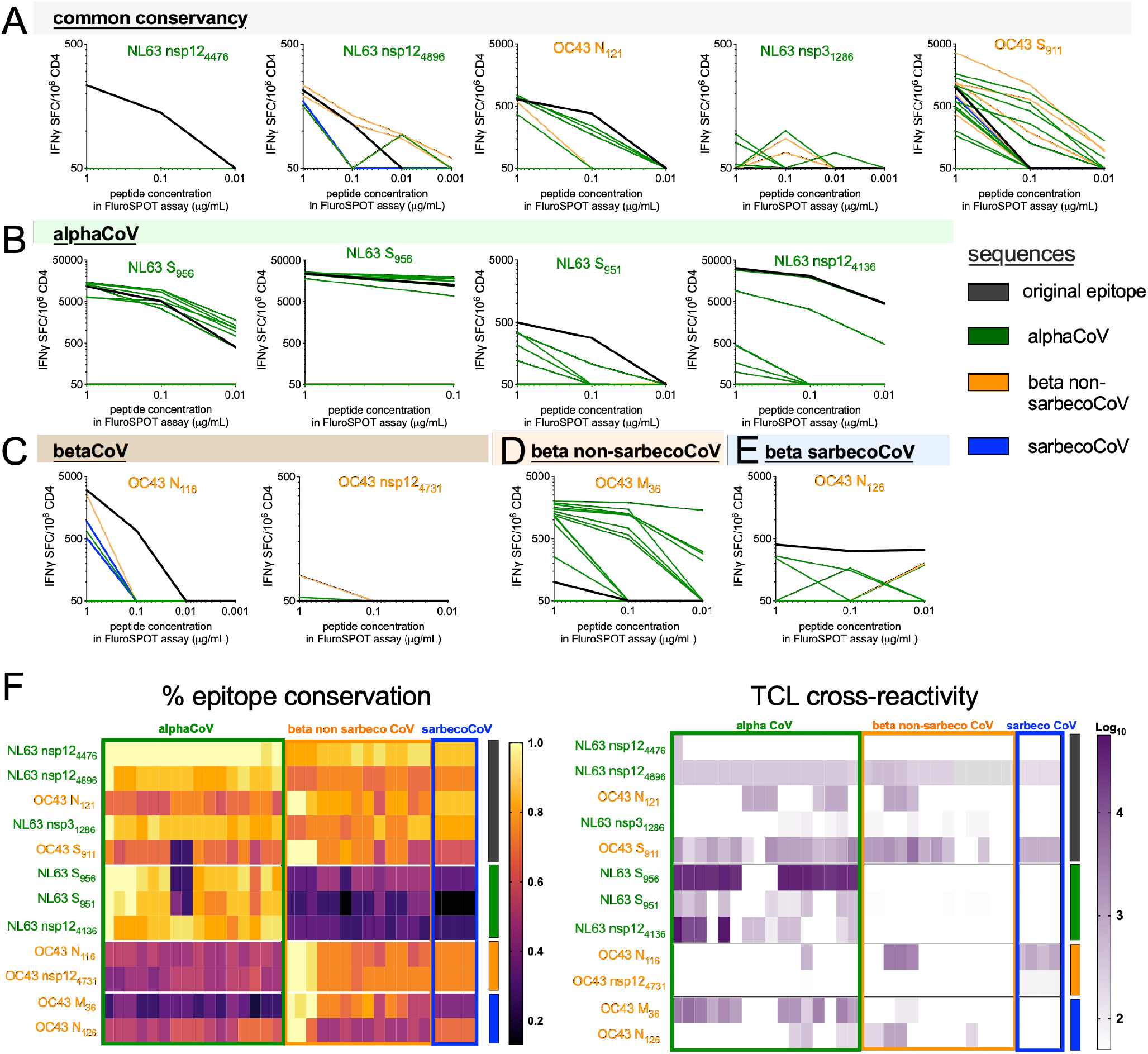
Cross-reactivity of NL63 and OC43 epitopes and homologous CoVs peptides. Twelve CCC epitopes were used to stimulate donor PBMCs and generate short-term T cell lines (TCL). The epitopes selected used specific NL63 or OC43 donor-epitope combinations based on the primary screen with the First Cohort. The TCLs are divided based on prediction of the original epitope selected and predicted on the basis of median sequence conservation >67% to be common (**A**) or specific for alphaCoV (**B**), betaCoV (**C**) and further segregating the betaCoV into non-sarbeco (**D**) and sarbeco (**E**) groups. After 14 days of *in vitro* expansion, each TCL was tested with the CCC epitope used for stimulation (black line in A, B, and C) and peptides corresponding to analogous sequences from other CoVs at three different concentrations (1, 0.1, and 0.01 μg/ml). IFNγ SFCs/10^6^ PBMCs are plotted for T cell lines stimulated with each peptide. Sequence identity and overall reactivity are shown as heatmaps in panel (**F**). The heatmap for the TCL reactivity represents the Log10 scale of the sum reactivity for the three concentrations of peptide tested.

The outcome of these experiments is also summarized in **Table 3** under the headings “First Cohort”. Overall, 7 out of 8 epitopes conserved in alphaCoVs showed T cell reactivity against alphaCoVs, and 6 out of 8 epitopes conserved in beta-non-sarbecoCoVs showed T cell reactivity for beta non-sarbecoCoVs. However, only 3 out of 8 epitopes conserved in beta-non-sarbecoCoVs showed T cell reactivity for sarbecoCoVs. In conclusion, the results show that cross-reactivity patterns are predicted by sequence conservation for alphaCoV and beta non-sarbecoCoVs. Conversely, cross-reactivity within sarbecoCoVsthat are phylogenetically more distant from other beta non-sarbecoCoVs was not well predicted by sequence conservation. Indeed,only SARS-CoV-2 was circulating in humans at the time the samples were obtained and all donors in this cohort were seronegative for SARS-CoV-2.

### Patterns of T cell cross-recognition of other Coronaviruses in an independent cohort

The results presented above show that while cross-reactivity for the alphaCoV and beta-non-sarbecoCoV groups could be predicted based on the known sequence conservation, sarbecoCoV reactivity was infrequent and not readily predicted based on sequence conservation. In the next round of experiments, we sought to verify these results in an independent cohort, and also investigate whether using alphaCoV, beta-non-sarbecoCoV or sarbecoCoV sequences for the *in vitro* peptide stimulation might modulate the cross-reactivity patterns. These experiments were performed with a new validation cohort (**Table 1**) as additional PBMCs from the previous cohort were largely not available. PBMC samples from the validation cohort were collected between October 2018 and August 2019 to ensure that donors had not been exposed to the sarbeco virus SARS-CoV-2 (**Table 1**). Preliminary experiments determined the *ex-vivo* reactivity in the validation cohort of previously identified epitopes. In these experiments, we tested both peptides corresponding to the OC43 and NL63 sequences and for 8 epitopes we detected reactivity to either the alphaCoV or betaCoV peptide version, or both (**Figure 4A-D**; left most part of the graphs).

**Figure 4.**
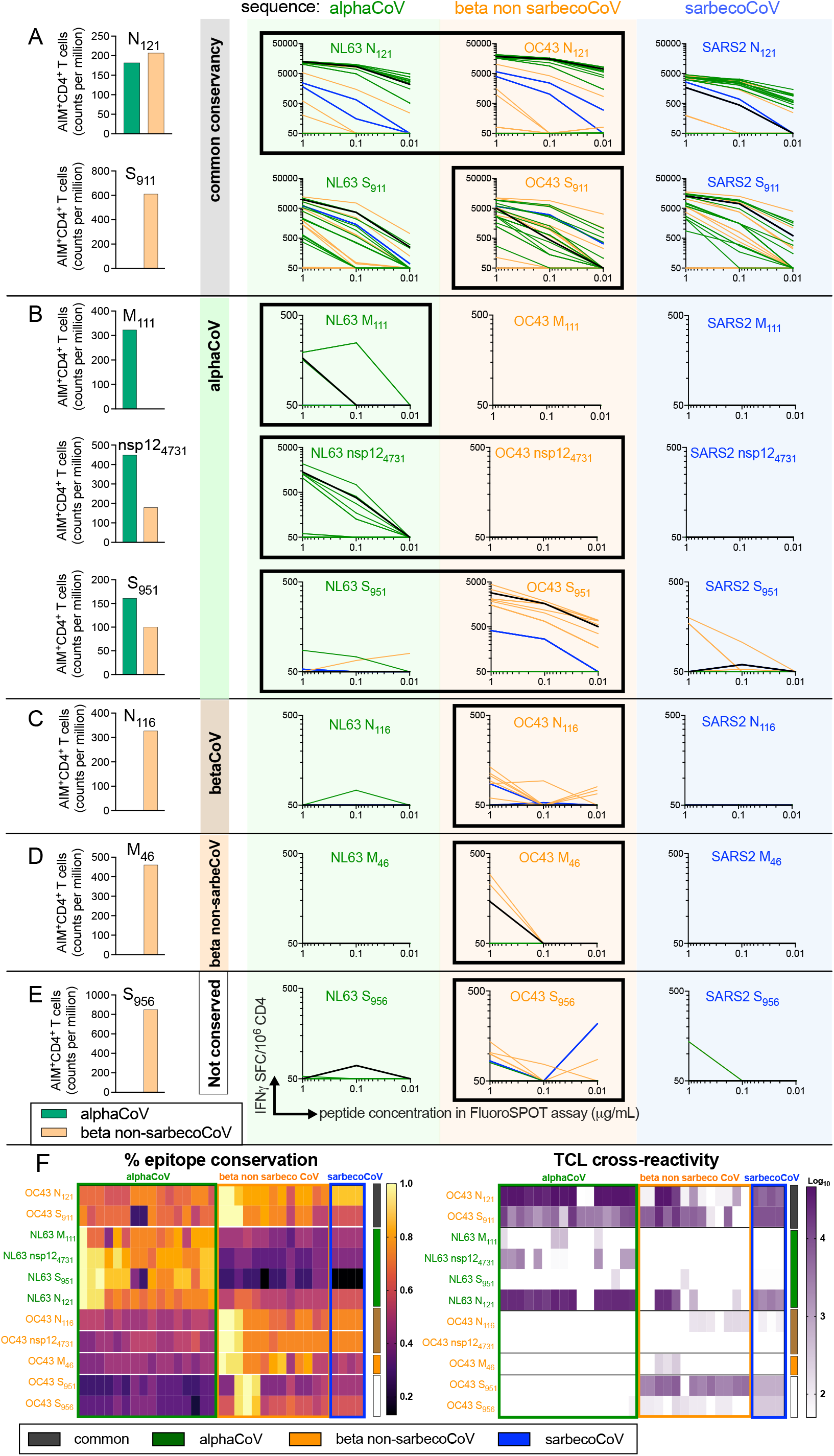
Cross-reactivity as a function of the initial peptide used for TCL generation. Shown here are 8 epitopes that had an *ex vivo* CCC response in the validation cohort of healthy donors (n=7). (**A-D**) The bar graphs show the T cell responses *ex vivo* to the NL63 (green) and OC43 (orange) epitopes as quantified by AIM assay. For each donor, 3 TCLs were generated based on NL63 (green), OC43 (orange), and SARS-CoV-2 (blue) prototype peptide sequences. After 14 days of *in vitro* expansion, each TCL was tested with the CCC epitope used for stimulation and peptides corresponding to analogous sequences from other CoVs at three different concentrations (1, 0.1 and 0.01 μg/ml). IFNγ SFCs/10^6^ PBMCs are plotted for TCL stimulated with each peptide. Common (**A**), alphaCoV-specific (**B**), betaCoV-specific (**C**), beta-non-sarbecoCoV-specific (**D**), and non-conserved (**E**) categories were selected based on predicted sequence identity. Sequence identity and overall reactivity are shown as heatmaps in panel (**F**). The heatmap for the TCL reactivity represents the Log_10_ scale of the sum reactivity for the three concentrations of peptide tested.

We then generated for each donor-peptide combination separate TCLs using the prototype NL63 or OC43 peptides, and then tested for cross-reactivity with the other alphaCoV and betaCoV sequences. We also used SARS-CoV-2 peptide sequences to generate TCL, and the results are described in the following section. The specific TCL reactivity for each epitope and virus sequence is shown in **Figure 4A-D**. Reactivity against alphaCoV sequences is shown in green, beta non-sarbecoCoV in orange, and sarbecoCoV in blue. The reactivity of epitopes conserved in all three groups is shown in **Fig. 4A**, alpha-specific (conserved only in alphaCoV) in **Fig. 4B**, beta-specific (conserved only in betaCoV) in **Fig. 4C**,beta non-sarbeco-specific in **Fig. 4D**,or not conserved in **Fig. 4E**. The results are also summarized as heatmaps in **Figure 4F**,depicting the epitope conservation across each viral species and the reactivity of each TCL against each peptide. The outcome of these experiments is also summarized in **Table 3** under the heading “Validation Cohort.”

We first analyzed the reactivity of TCLs generated by *in vitro* stimulation with the same peptide epitopes for which *ex vivo* reactivity was detected in that specific donor; these instances are highlighted by a black highlighted margin around the graphs in **Fig 4**. In those instances, in the case of epitopes that were predicted to be conserved across CoVs, 2 out of 2 instances showed cross-reactivity for alphaCoV, beta non-sarbecoCoV and sarbecoCoV when the homologous epitope was used for the TCL generation (**Fig. 4A**). 3 epitopes with sequence conservation within alphaCoV, had *ex vivo* reactivity with the alphaCoV NL63 sequence and also showed cross-reactivity to other alphaCoV sequences after TCL expansion (**Fig. 4B**). The NL63 M_111_ epitope is alpha-specific at the level of conservation and the TCL was associated with a predominant alpha-specific reactivity (**Fig. 4B**).

In the case of nsp12_4731_ and S_951_, *ex vivo* reactivity was detected for both the NL63 and OC43 epitopes, potentially reflective of either multiple exposures and/or cross-reactivity. For nsp124731, the NL63 sequence is conserved within alpha but not beta (alphaCoV= 0.8; beta non-sarbecoCoV= 0.33), and conversely the OC43 sequence is conserved within beta but not alpha (alphaCoV= 0.47; betaCoV= 0.73) (**Table 3**). The TCL obtained by NL63 *in vitro* stimulation was alpha specific, while no *in vitro* expansion was noted in the case of the TCL stimulated with the OC43 sequence (**Fig. 4B**). In the case of S_951_, the TCL expanded with the OC43 epitope was mostly beta reactive, and conversely, the TCL expanded with the NL63 epitope was alpha-reactive (**Fig. 4B**). These results are consistent with the fact that the NL63 S_951_ epitope is well conserved within alpha but not beta (alphaCoV= 0.87; beta non-sarbecoCoV= 0.27) (**Table 3**). In contrast, the OC43 S_951_ epitope is not well conserved within alpha or beta (alphaCoV= 0.33; beta non-sarbecoCoV= 0.4) (**Table 3**), however the beta-specific cross-reactivity could be attributed to the subset of betaCoV sequences that are conserved for this epitope, as seen in **Fig. 4F**.

The betaCoV-specific OC43 N_116_ epitope induced beta-specific cross-reactivity as expected, based on sequence conservation and *ex vivo* reactivity (beta-non-sarbeCoV= 0.73; sarbecoCoV= 0.73; **Fig. 4C** and **Table 3**). The beta-non-sarbeco conserved epitope OC43 M_46_ was associated with OC43 reactivity *ex vivo*, and the associated TCL displayed predominant beta-non-sarbeco reactivity (**Fig. 4D** and **Table 3**). The S_956_ epitope was associated with OC43 beta reactivity in the specific donor tested. Consistent with this observation, the TCL maintained this pattern of prevalent beta reactivity (**Fig. 4E**). This result is consistent with the fact that the OC43 S_956_ epitope is well conserved within some beta but not alpha (alphaCoV= 0.4; beta non-sarbecoCoV=0.5) (**Table 3**).

### Overall efficacy of sequence conservation and pre-existing reactivity as a predictor of coronavirus cross-reactivity

The overall data from the first and validation cohorts combined was evaluated for parameters that might guide selection of epitopes linked with cross-reactive T cell responses. **Table 4** details how frequently sequence conservation of 67%, or above, was associated with experimentally verified cross-reactivity in each of the taxonomic coronavirus groups. When the three groups are considered together, the 67% median sequence conservation threshold predicts T cell cross-reactivity in 73% of the cases (considering 13+10+6=29 instances of crossreactivity/ 15+13+12=40 tested). When only alphaCoV and beta non-sarbecoCoV are considered, cross-reactivity is correctly predicted in 89% of cases, while in only betaCoV (Beta non-sarbecoCoV and sarbecoCoV) the prediction accuracy is 69%. When we look within each group separately, the predictive capacity is 87% and 77% if only alpha or beta non-sarbecoCoV groups are considered. This is in contrast with the fact that experimental T cell cross-reactivity was observed only in 50% of the cases of sequences conserved within sarbecoCoV (**Table 4**).

**Table 4.** Summary of sequence conservation and TCL cross-reactivity as predictor of CoV cross-reactivity.

|  | Conserved<br>(>67%) | Cross-reactive | %<br>correct |
| --- | --- | --- | --- |
| Alpha | 14 | 13 | 93% |
| Beta non-sarbeco | 14 | 12 | 86% |
| Sarbeco | 12 | 6 | 50% |
| All | 40 | 29 | 73% |
| Alpha+Beta non-sarbeco | 28 | 25 | 89% |
| Beta non-sarbeco+sarbeco | 26 | 18 | 69% |

In conclusion, the results show that cross-reactivity patterns are predicted by sequence conservation for alphaCoV and beta non-sarbecoCoVs. Conversely, cross-reactivity with sarbecoCoVs (which were not circulating in humans at the time the samples were obtained) was not well predicted by sequence conservation. Relevant to the hypothesis that pre-exposure might influence the capacity of experimentally observing T cell cross-reactivity, we examined the association between *ex vivo* reactivity and measured cross-reactivity in the expanded TCLs. Indeed, when we consider the OC43 and NL63 epitopes in **Fig. 4**, for which *ex vivo* reactivity was experimentally determined, *ex vivo* reactivity was detected in 11 instances, of which 9 showed cross-reactivity in the expanded TCLs. Of the 5 epitopes for which *ex vivo* reactivity was not detected prior to *in vitro* expansion to derive specific TCLs, only 1 yielded cross-reactive TCLs (p=0.0357 by the Fisher exact test). This observation demonstrates a correlation of *ex-vivo* reactivity with cross-reactivity after TCL expansion.

As stated above, in parallel experiments we also used SARS-CoV-2 peptide sequences for re-stimulation, to examine whether this approach could increase the frequency and extent of CCC cross-reactivity within sarbecoCoV sequences. The results are shown in **Figure 4** and summarized in **Table 3**.

Overall, when either NL63 or OC43 sequences were used for re-stimulation, sarbecoCoV cross-reactivity was noted in 3/12 instances for the First Cohort (25%), and in 7/16 (44%) for the Validation Cohort, for an overall frequency of 10/28 (36%). When the SARS-CoV-2 sequences were utilized in the TCL generation, sarbeco cross-reactivity was noted in 2/8 instances (25%). Thus, the use of SARS-CoV-2 sequences to expand TLCs was not associated with an increase in cross-reactivity, and was actually associated with a trend toward lower reactivity. This might reflect the possibility that stimulation with CCC epitopes might be most related to the original *in vivo* immunogen in this pre-pandemic cohort studied, and thereby also a more effective *in vitro* stimulus for TLC expansion.

## DISCUSSION

The data from the present study addresses three main issues-what is the pattern of antigens recognized by human CD4^+^ memory T cells recognizing CCC, what is the relation of the associated epitope repertoire with the SARS-CoV-2-associated epitope repertoire, and to what extent can cross-reactive T cell responses between different taxonomic CoV groups can be observed and predicted. In addition, the studies revealed a prominent role of pre-existing immunity as a driver of development of cross-reactive T cell responses.

In terms of the antigens recognized as immunodominant by T cell responses, this is the first report systematically evaluating which antigens are recognized by human memory CD4^+^ T cell responses in alpha and beta CCC. NL63 and OC43 immunodominant proteins included the structural proteins S, M, and N, and the non-structural protein nsp3. This pattern of immunodominance is similar to what was previously observed in the context of SARS-CoV-2, suggesting these antigens are common targets across multiple CoVs ^39,49,52^. This is in line with the work of others who described immunodominant CD4^+^ T cell responses to CCC-conserved epitopes ^35–38,45,55–61^, which have been previously reviewed ^30,62^. Thus, the dominance of these antigens in coronavirus recognition might reflect conserved and common mechanisms, such as high levels of expression ^33^.

Our study also revealed interesting features of immunodominance specific to CCC. NL63-specific T cell responses prominently recognized nsp2, and OC43-specific T cells recognized nsp12, previously described in the SARS-CoV-2 context ^49,52^ as recognized by cross-reactive T cells in the context of asymptomatic SARS-CoV-2 infection in healthcare workers ^63^. Differences in the profile of antigens recognized by different coronaviruses is also to be expected given that some antigens are specifically encoded in some but not other coronavirus genomes, such as the NL63 NP3 protein, which is not found in OC43 or SARS-CoV-2 and the OC43 NS2a protein, which is not encoded in NL63 or SARS-CoV-2 (see Methods for virus sequence information). While these antigens could perhaps have some diagnostic value, our data suggest that they are relatively minor targets for human T cells. Conversely, the fact that certain antigens are broadly conserved targets for human T cell recognition support the notion that these antigens could be utilized to elicit broadly reactive responses.

The present study also provides the first account of the epitope repertoire associated with human CCC-specific memory CD4^+^ T cells. We found an average breadth of 7 epitopes (range 1 to 24) per donor being recognized. This is 2-3 fold fewer than what we previously observed in the case of SARS-CoV-2, where an average of 19 CD4 epitopes/donor (range 1 to 63) were identified using a similar experimental strategy ^52^. The lower number of epitopes detected for the CCC is likely a reflection of the fact that the SARS-CoV-2 epitope identification studies were performed 2 months after SARS-CoV-2 infection ^52^, while the present CCC studies were performed in subjects of unknown and presumably less recent exposure. The identified epitopes were associated with predicted promiscuous binding capacity to a panel of frequent HLA alleles, confirming data obtained in several other systems ^64–67^ and utilized to develop algorithms to predict dominant CD4^+^ T cell epitopes ^68–70^.

The epitope repertoire identified by *ex vivo* reactivity in NL63 and OC43 encompassed a total of 165 epitopes. These epitopes were largely undescribed, with only 10 epitopes overlapping with epitopes previously described in the IEDB ^71^, even by allowing a rather loose criteria of 67% sequence homology. The overlap between the repertoire of epitopes recognized in NL63 and OC43 in the current study, as also defined by a 67% sequence homology or more, was limited to 2 epitopes. Furthermore, of relevance to the issue of T cell cross-reactivity across different coronaviruses, discussed below, the NL63 and OC43 epitope repertoire was largely non-overlapping (6 epitopes; 4% overlap) with the repertoire of SARS-CoV-2 specific CD4^+^ T cell responses previously described ^52^. This is consistent with previous observations that described pre-existing cross-reactive memory SARS-CoV-2 responses from unexposed individuals that share sequence homology with CCC ^33,36,39^, but also that the T cell repertoire that develops upon SARS-CoV-2 infection, is largely non-overlapping with the repertoire recognized by pre-existing cross-reactive memory SARS-CoV-2 responses from unexposed individuals ^52^.As discussed above in the Introduction, several lines of evidence suggest that T cells play a role in limiting disease severity and terminating infection, in the context of SARS-CoV-2 infection ^7,9–11^. The preservation of T cell reactivity against variants and the impairment of neutralizing antibodies correlates with preservation of protection against severe disease and decreased protection from infection ^16,18,19,72^. Directly applicable to the present study are the observations, also summarized in the Introduction, that pre-existing cross-reactive immunity associated with CCC are beneficial in the context of SARS-CoV-2 infection and vaccination ^31,32,63,73^. Based on this rationale it has been proposed that immunodominant T cell regions conserved across CoVs may be of interest in the context of inducing a panCoV T cell response ^1^.

What strategies can be utilized to predict and detect such cross-reactive responses? Bioinformatic analysis of sequence conservation in panels of different viral species was shown to be effective in informing selection of potential cross-reactive T cell epitopes ^33,36,70^. Crossreactive epitopes were associated with overall 67% or greater sequence conservation, in agreement with previous studies in the context of SARS-CoV-2 and other viral infections such as ZIKV and VZV ^36,58,74,75^. Our results provide the largest data set available to address this issue, with cross-reactivity data involving 18 different epitopes and 37 TCLs, testing a total of 594 different viral variant epitope sequences. Within the alpha and beta-non-sarbeco groups, the degree of sequence conservation was frequently reflected in CD4^+^ T cell cross-reactivity. Across alpha and betaCoV groups, we correctly predicted cross-reactivity in 29 out of 40 CoV-conserved epitopes considered (73% of the cases). This result validates the use of the 67% sequence identity value to predict T cell cross-reactivity. Previous studies on different viral species reported conserved T cell epitopes were also observed. In the influenza system, CD4^+^and CD8^+^ T cells epitopes conservation in multiple influenza strains were reported ^26,76^. In the context of flaviviruses, despite an overall low degree of cross-reactivity between different flaviviruses ^77^, such as Dengue, Yellow Fever, and Zika, it has been shown that T cell epitopes conserved among these viruses can protect against disease in animal models ^78,79^. Overall, cross-reactivity between viruses is more often observed across viruses with closer phylogenetics relations and greater sequence homology ^40,44^.

A remarkably different situation was observed when the cross-reactivity with sarbecoCoV was considered. In this case, there were few instances of T cell cross-reactivity, even in cases with higher sequence conservation, and a total of 6 out 12 instances of cross-reactivity was observed, corresponding to 50% of cases. The most straightforward explanation of this result in pre-pandemic or SARS-CoV-2-seronegative samples is that previous exposure to alpha and beta-non-sarbeco viruses is an important determinant of cross-reactivity alongside the degree of conservation. Indeed, sarbecoCoV are more phylogenetically distant and more different from other betaCoV including OC43. Accordinglythe lack of previous exposure to sarbecoCoVs in these samples, being collected pre-pandemic, was associated with lower degree of crossreactivity amongst different representative of sarbecoCoV sequences including SARS-CoV-2 and even if those sequences were relatively conserved. Hence, the lower cross reactivity observed in this study fits with the lack of previous exposure to sarbecoCoVs and suggest this is an important determinant of crossreactivity alongside the degree of conservation.We then asked if this bias in cross-reactivity using the *in vitro* expansion was due to the fact that we used only OC43 or NL63 epitopes and accordingly we simultaneously stimulated the same donor with the three peptide variants and we found that the cross-reactivity was not increased by using SARS-CoV-2 sequences.

Our overall goal is to be able to generate and amplify CD4^+^ T cells broadly cross-reactive with sarbecoCoVs of zoonotic origin and of potential pandemic concern.. Our data suggest that exposure is a contributing factor, therefore, future studies should focus on immunity following SARS-CoV-2 exposure. The large numbers of individuals that have been exposed to SARS-CoV-2 world-wide suggests that this should be a feasible goal. Selecting SARS-CoV-2 epitopes and antigenic regions associated with high immunodominance, indicating their recognition in the general human population, and broad conservation amongst other sarbecoCoVs and possibly other beta and alpha coronaviruses, will give the best opportunity of expanding and eliciting cross-CD4^+^ T cells broadly reactive for a number of different CoV species.

### Limitations and future directions

One limitation of this study is that the time of the most recent CoV exposure of the donors tested is unknown. We hypothesize that some of the reactivity we observed was impacted by the most recent CoV exposure. Additionally, our study did not evaluate cross-reactivity at the level of CD8^+^ T cells, and in SARS-CoV-2 exposed or vaccinated subjects. An additional limitation is that we did not perform serology tests to check for exposure to other zoonotic CoV species that may contribute to T cell cross-reactivity, however, exposure to other zoonotic CoVs has not been reported in the area of sample collection and it is reasonable to assume no or minor impact on the study’s results.

## ACKNOWLEDGEMENTS

This study has been funded by the NIH/NIAD P01 AI168347 to A.G, Contract No. 75N93019C00065 to A.S and D.W., and HHS75N93019C00076 to R.H.S. A.T. was supported by a PhD student fellowship through the Clinical and Experimental Immunology Course at the University of Genoa, Italy. We thank Gina Levi and the LJI clinical core for assistance in sample coordination and blood processing.

## AUTHOR CONTRIBUTIONS

Conceptualization: A.G., R.H.S. and A.S.; Data curation and bioinformatic analysis, Y.Z.; Formal analysis: A.T., N.M., and T.M.N; Funding acquisition: A.S. and R.H.S.; Investigation: A.T., Y.Z., N.M., T.M.N., R.H.S., A.S and A.G.; Project administration: A.F. Resources: D.W., S.M., E.P., J.D. and A.S.; Supervision: G.F., L.P., R.H.S., A.S., and A.G.; Writing: A.T., Y.Z., R.H.S., A.S., and A.G.

## DECLARATION OF INTEREST

A.S. is a consultant for Gritstone Bio, Flow Pharma, Moderna, AstraZeneca, Qiagen, Fortress, Gilead, Sanofi, Merck, RiverVest, MedaCorp, Turnstone, NA Vaccine Institute, Emervax, Gerson Lehrman Group and Guggenheim. LJI has filed for patent protection for various aspects of T cell epitope and vaccine design work.

## STAR METHODS

### RESOURCE AVAILABILITY

#### Lead Contact

Further information and requests for resources and reagents should be directed to the lead contact, Dr. Alba Grifoni, (A.G.).

#### Materials Availability

Epitope pools used in this study will be made available to the scientific community upon request, and following execution of a material transfer agreement, by contacting Dr. Alba Grifoni and Dr. Alessandro Sette.

#### Data and Code Availability

The published article includes all data generated or analyzed during this study, and summarized in the accompanying tables, figures and supplemental materials.

## EXPERIMENTAL MODEL AND SUBJECT DETAILS

### Human Subjects

Blood samples from healthy adult donors were obtained from the San Diego Blood Bank (SDBB). Subjects were considered eligible for this study if they fulfilled the SDBB criteria to donate blood and if they were tested and found negative for SARS-CoV-2 RBD IgG serology. Overview of the cohort analyzed is summarized in **Table 1**. Whole blood was collected from all donors in heparin coated blood bags and processed as previously described (Tarke et al., 2021). Briefly, peripheral blood mononuclear cells (PBMC) were isolated by density-gradient sedimentation using Ficoll-Paque (Lymphoprep, and 90% heat-inactivated fetal bovine serum (FBS; Hyclone Laboratories, Logan, UT) and stored in liquid nitrogen until used in the assays. Each sample was HLA typed by Murdoch University in Western Australia, an ASHI-accredited laboratory. Typing was performed for the class II DRB1, DQB1, and DPB1 loci.

## METHOD DETAILS

### Peptide Pools

*Preparation of 15-mer peptides and subsequent megapools and mesopools.* To identify CCC-specific T cell epitopes, we synthesized 15-mer peptides overlapping by 10 amino acids and spanning the entire NL63 and OC43 proteomes. All peptides were synthesized as crude material (TC Lab, San Diego, CA) and individually resuspended in dimethyl sulfoxide (DMSO) at a concentration of 20 mg/mL. Aliquots of peptides were either pooled by antigen (megapools; MP) or by ten peptides each (mesopools). The MP required an additional step of sequential lyophilization as previously reported (Carrasco Pro et al., 2015). MPs were resuspended at 1 mg/mL in DMSO, while mesopools were resuspended at 2mg/mL.

### SARS-CoV-2 ELISA

The SARS-CoV-2 RBD ELISA has been described in detail elsewhere (Dan et al., 2021; Grifoni et al., 2020a). Briefly, 96-well half-area plates (ThermoFisher 3690) were coated with 1 μg/mL SARS-CoV-2 Spike (S) Receptor Binding Domain (RBD) and incubated at 4°C overnight. On the following day plates were blocked at room temperature for 2 hours with 3% milk in phosphate buffered saline (PBS) containing 0.05% Tween-20. Then, heat-inactivated plasma was added to the plates for another 90-minute incubation at room temperature followed by incubation with conjugated secondary antibody, detection, and subsequent data analysis by reading the plates on Spectramax Plate Reader at 450 nm using SoftMax Pro.

### CCC ELISA

OC43 and NL63 RBD ELISA were carried out as previously described ^80,81^. Briefly, coating was performed by Streptavidin (Invitrogen) at 4 μg/mL in Tris-Buffered Saline (TBS) pH 7.4 for 1 h at 37°C followed by blocking with Non-Animal Protein-BLOCKER^™^ (GBiosciences). Then biotinylated spike RBD antigens for OC43 and NL63 were added at 1 μg/mL at 37°C for 1 h. All plasma samples were heat-inactivated before usage to minimize risk of residual virus in serum and then incubated at serial dilution followed by multiple washes and incubation with horseradish peroxidase-conjugated secondary Goat Anti-Human secondary IgG (Cat No: 109-035-008, Jackson ImmunoResearch) at 1:40,000 dilution in 3% milk at 37°C for 1 h. The resulting plate was washed and 3,3’,5,5’ -Tetramethylbenzidine (TMB) Liquid Substrate (Sigma-Aldrich) was added for optical density (OD) measurement at 405 nm after stopping the reaction with 50 μl of 1 N HCl.

### Flow Cytometry

*Activation induced cell marker (AIM) assay.* The AIM assay for epitope identification was performed mirroring the previously described protocol (Tarke et al., 2021). Cryopreserved PBMCs were thawed in RPMI 1640 media supplemented with 5% human AB serum (Gemini Bioproducts) in the presence of benzonase [20 μl/10 ml]. Cells were stimulated for 24 hours in the presence of CCC specific MPs or mesopools at 1 μg/ml and then deconvoluted with 15-mer peptides [10 μg/ml] to reach the epitope level. Stimulation was carried out in 96-wells U bottom plates with 1×10^6^ PBMC per well. An equimolar amount of DMSO was used as negative control in triplicates, while stimulation with phytohemagglutinin (PHA, Roche, 1 μg/ml) was included as the positive control. The cells were stained with CD3 AF700 (2:100; Life Technologies Cat# 56-0038-42), CD4 BV605 (1:100; BD Biosciences Cat# 562658), CD8 BUV496 (2:100; Biolegend Cat#612942), CD14 V500 (2:100; BD Biosciences Cat# 561391), CD19 V500 (2:100; BD Biosciences Cat#561121), and Live/Dead eFluor 506 (25:1000; eBioscience Cat# 65-0866-18). Activation was measured by the following markers: CD137 APC (4:100; Biolegend Cat# 309810) and OX40 PE-Cy7 (2:100; Biolegend Cat#350012). All samples were acquired on ZE5 cell analyzer (Bio-rad laboratories) and analyzed with FlowJo software (Tree Star).

### *In vitro* expansion of OC43 and NL63 specific T cells lines (TCLs) and cross-reactivity assessment by FluoroSPOT assays

*In vitro* expansion of OC43 and NL63 specific T cells was carried out for 14 days to generate epitope-specific T Cell Lines (TCLs). The TCLs were set up using donors selected from the NL63 and OC43 epitope identification screening. The PBMCs were expanded using specific epitope/donor [1 μg/ml] combinations chosen on the basis of the NL63 and OC43 CD4^+^ T cell epitope screening. IL-2 was added on day 3, 7, and 11. On the 14th day, the cells were harvested and triplicates of 5×10^4^ PBMCs were incubated in the presence of the epitope used for expansion and subsequent homologous CoV peptides based on representatives’ sequence selection. Each peptide was tested at 2 to 5 different serial concentrations depending on cells availability after the 14 days of culture (1 μg/mL, 0.1 μg/mL, 0.01 μg/mL, 0.001 μg/mL, and 0.0001 μg/mL) and measured by IFNγ FluoroSPOT assay as previously reported ^36^. Briefly, cells were incubated in presence of peptide stimulation for 20 hours at 37 C, 5% CO_2_ at a concentration of 1×10^5^ cells/mL. Cells were then incubated with IFNγ mAb (7-B6-1-BAM Mabtech, Stockholm, Sweden) for 2 hours and developed.

## BIOINFORMATIC AND STATISTICAL ANALYSIS

### Sequence download and quality control

The genome and protein sequences used in this study were downloaded from the Virus Pathogen Resource (ViPR; https://www.viprbrc.org/) and Bacterial and Viral Bioinformatics Resource Center (BV-BRC; https://www.bv-brc.org)^82^ websites on June 17, 2021. Given the unprecedented number of SARS-CoV-2 genomes, to make the data size more manageable, the SARS-CoV-2 reference dataset (1438 strains as of June 17, 2021) computed by the ViPR team was used. For alphacoronavirus and non-SARS-CoV-2 betacoronavirus, all available sequences in ViPR were used. Potential laboratory strains and low-quality sequences were filtered out using custom scripts.

### Representative virus selection

Representative virus selection is summarized in **Figure S3**. To select viruses that are representative of each taxon group (alphacoronavirus, non-sarbeco betacoronavirus, and sarbecovirus), a targeted sampling approach was used which leverages sequence identity, sequence and annotation quality, host, isolation date and region, RefSeq designation, and phylogenetic structures. First, all genome sequences were clustered based on sequence identity using cd-hit with the non-greedy option that assigns shorter sequences to the closest cluster. To get the desired number of representative viruses, a 0.80 identity threshold was used for the alpha and non-sarbeco beta groups, while a 0.999 threshold was used for the sarbeco group as sarbeco strains have very high similarity in sequence identity. Second, the taxa and metadata (host, isolation country, and isolation year) associated with the sequences in each cluster were extracted using custom scripts. Third, target clusters were selected based on the cluster size (at least 2 for alpha and non-sarbeco beta groups, and at least 10 for the sarbeco group) and host (human, bat, camel, hedgehog, or pangolin). Fourth, representative viruses were selected for each target cluster. Specifically, if a cluster contains NCBI RefSeq sequences (1 or 2 sequences), the RefSeqs were selected as the representatives. Otherwise, a virus with complete metadata, good quality protein annotations and from a recent subcluster was selected as the representative. Finally, since the sarbeco group has a much narrower taxonomic scope, an extra iteration of phylogeny-based sampling was performed to ensure the best diversity representation. Using the genome sequences of the selected sarbeco candidates, a phylogenetic tree was built and visualized on the ViPR website. Representatives were selected to cover the major phylogenetic clusters. This targeted sampling process resulted in the selection of 16 alpha, 13 non-sarbeco beta and 4 sarbecoviruses as representatives.

### T cell epitope homolog identification

To find the epitope homologs in the representative viruses, the epitope homologous region in each representative was identified and then the optimal k-mer was found in this region (**Figure S4**). Specifically, each epitope was mapped to the virus taxon’s RefSeq protein. Then the RefSeq protein sequence harboring the mapped epitope was aligned with each taxon group’s protein sequences using the mafft program einsi mode. In the resulting alignment, the epitope mapped peptide in each virus was defined as the seed. Using the seed, the search space for finding the optimal k-mer, where k is the length of the epitope, was defined. If the seed was k or longer, the search space was the seed itself. Otherwise, the search space was expanded to include (k – seed_length) additional residues on both sides of the seed, unless the boundary of the protein was reached:

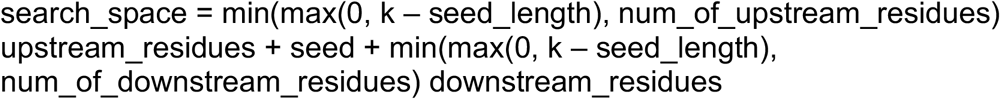

To find the optimal k-mer, each k-mer in the search space was calculated with an identity score, and the k-mer with the maximum score was selected. In case of ties, the leftmost k-mer with the maximum score was selected:

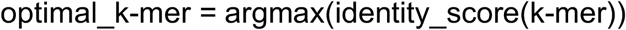

The k-mer identity score was defined as the maximum number of matched residues in all possible non-gap alignments divided by the epitope length k:

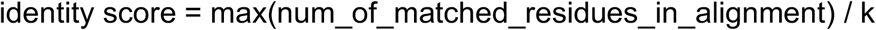

A non-gap alignment was defined as a pairwise alignment of the k-mer and the input epitope that only allowed shifting and substitutions, but no internal indels. Each k-mer had 2 * k - 1 such alignments.

## SUPPLEMENTARY MATERIALS

### SUPPLEMENTARY FIGURE LEGENDS

**Figure S1.**
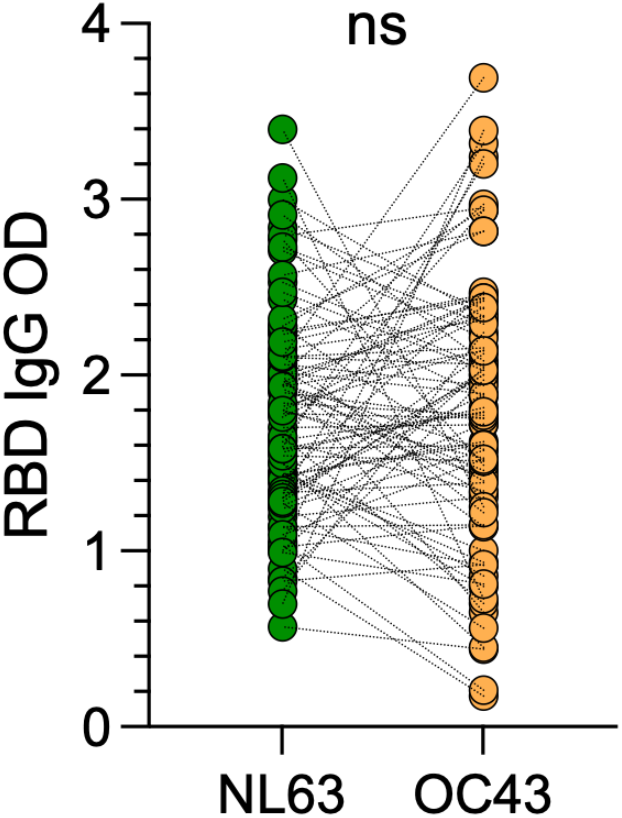
NL63 and OC43 serology of the cohort of healthy donors. The RBD IgG OD for NL63 and OC43 are shown for the healthy donors (n=88) in this study and the dotted lines connect the same donor analyzed for NL63 or OC43. Comparison of serum antibodies to NL63 and OC43 RBD was performed using the Wilcoxon rank-sum test (also see **Table 1**).

**Figure S2.**
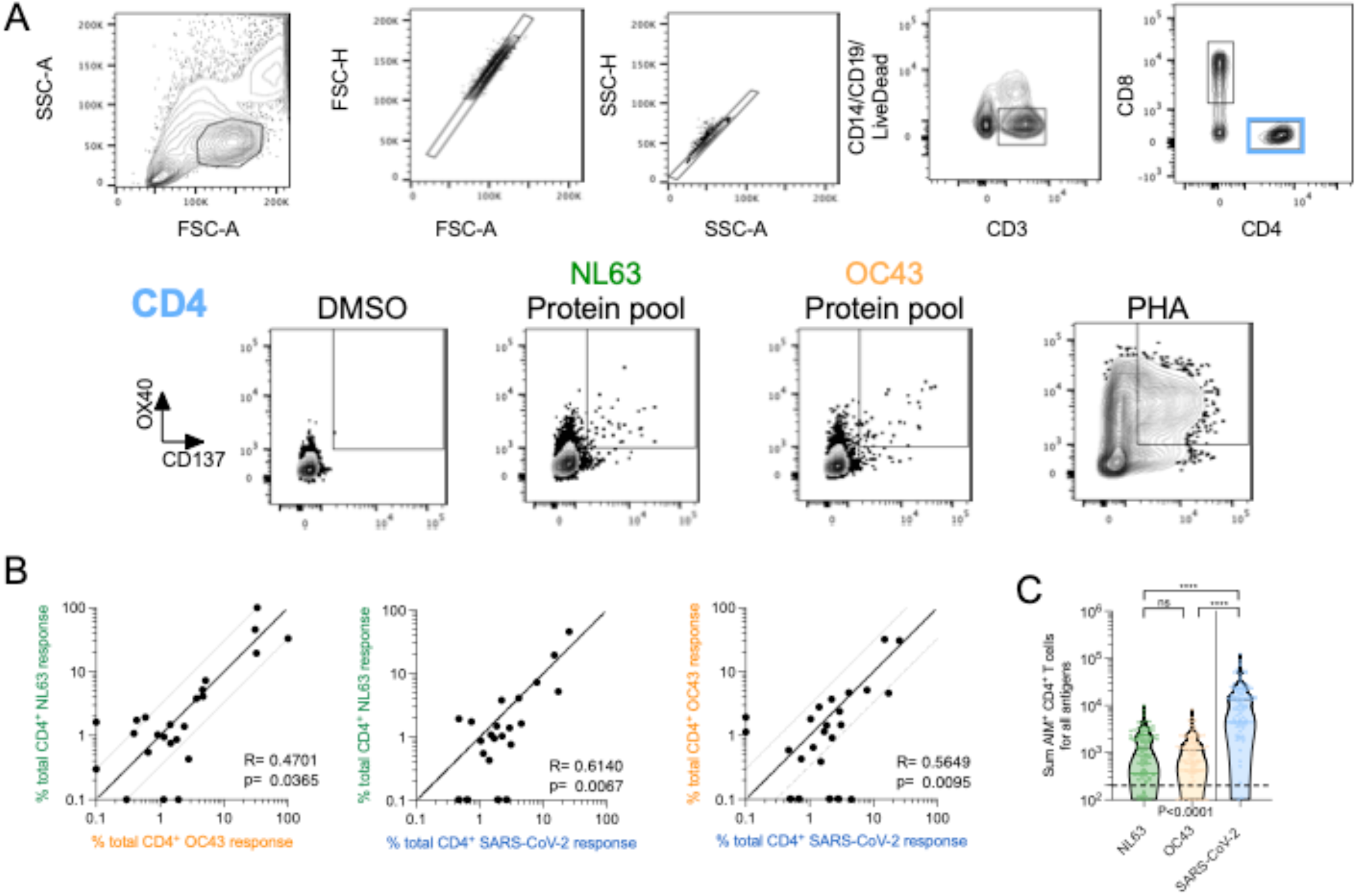
Total AIM^+^ CD4^+^ T cell reactivity against antigens related to NL63, OC43 and SARS-CoV-2. **A**)The gating strategy for the AIM assay is shown. **B**) Data are expressed as sum counts of OX40^+^ CD137^+^ CD4^+^ T cells for each individual positive antigen for NL63 (green), OC43 (orange) and SARS-CoV-2 (blue). Historical data on T cell responses to SARS-CoV-2 is from COVID-19 convalescent donors (n=99) originally published in Tarke et al. ^52^. OC43 and NL63 T cell reactivity reflects the First Cohort of healthy donors (n=88) described in this study. Pairwise correlation among the three viruses per protein is shown together with Spearman correlation R and p-value. **C**) Comparison of the three viruses are shown. Data are compared by Kruskal Wallis test (P<0.0001, below) as well as Mann-Whitney (above) for each of the paired comparisons. **** p < 0.0001. Refers to **Table 1** and **Figure 1**.

**Figure S3.**
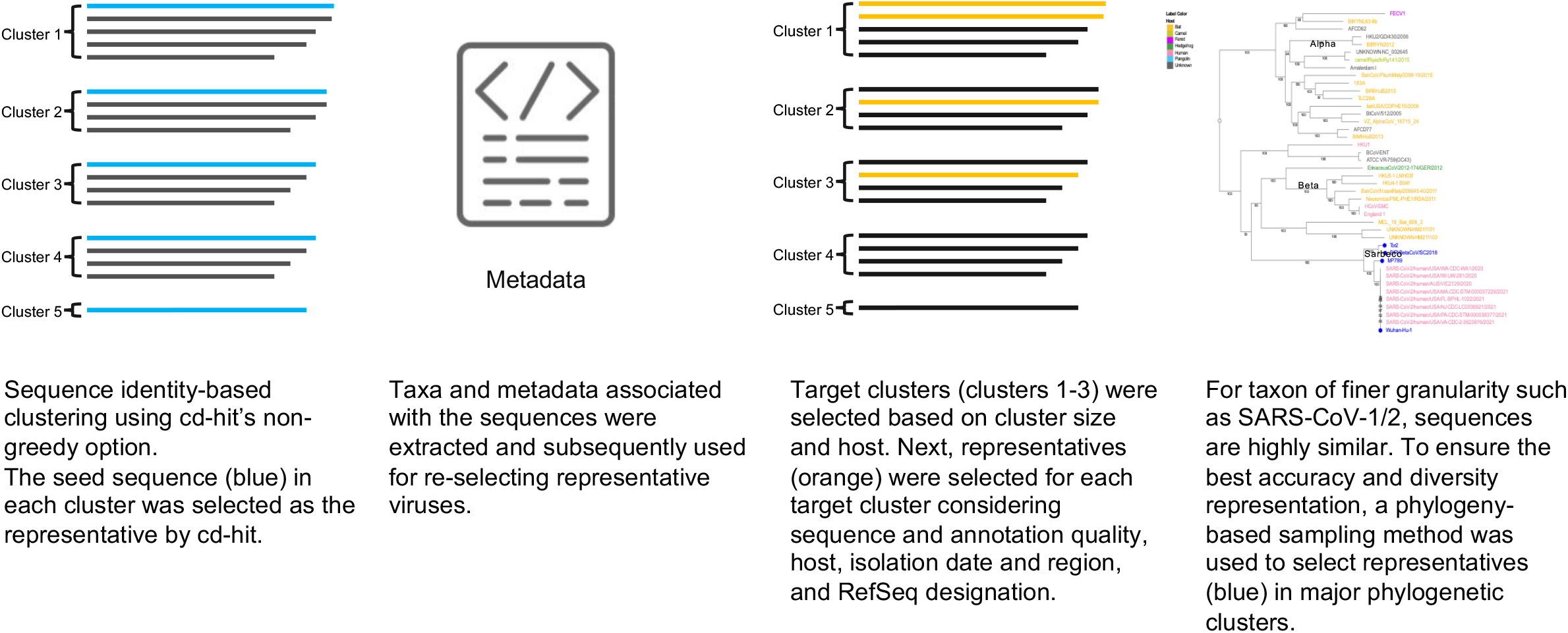
Representative virus selection. To select representative viruses for each taxon group (alphaCoV, sarbeco, non-sarbeco betaCoV), a targeted sampling approach was used which leverages sequence identity, sequence and annotation quality, host, isolation date and region, RefSeq designation, and phylogenetic structure (see Methods and **Table 2** for details).

**Figure S4.**
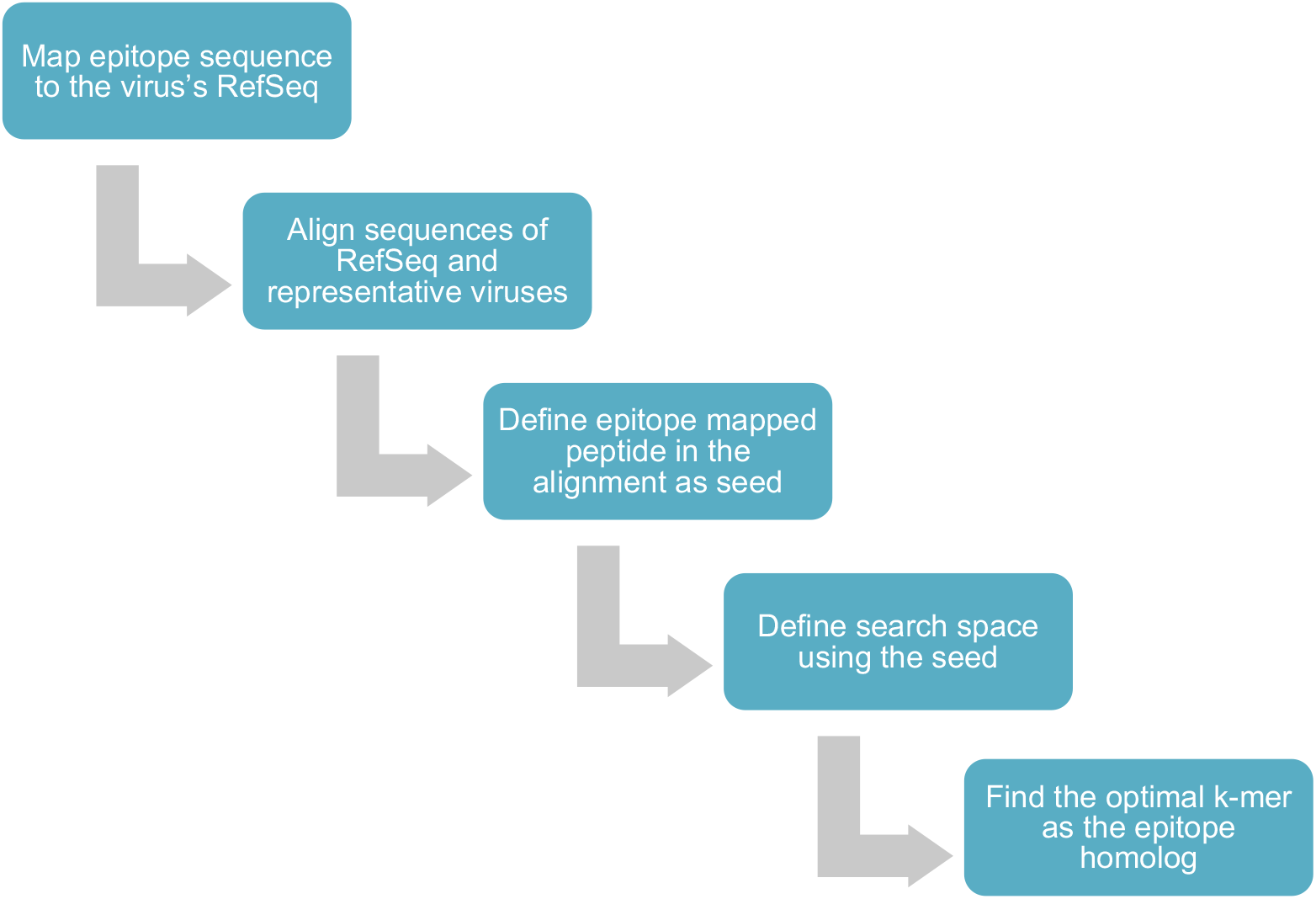
Pipeline to establish the degree of sequence conservation of each epitope. Epitopes were mapped to each of the representative viruses to identify the epitope homologs using an alignment-based, k-mer finding approach (see Methods for details). This process first identified the epitope homologous region in each representative and then selected the optimal k-mer in the region as the epitope homolog. The level of sequence conservation of the homologous epitope regions is shown in **Figure 2**.

## SUPPLEMENTARY TABLE LEGENDS

**Table S1. HLA typing of donor cohort.**

**Table S2. List of CD4^+^ T cell epitopes identified in this study.** NL63 and OC43 epitopes information pertaining protein composition and location are included togther with the sequence identity values related to each of the representative sequence for Sarbecoviruses (n=10), Betacoronaviruses excluding Sarbecoviruses (n=15) and Alphacoronaviruses (n=15).

